# Alone in a crowd: Effect of a nonfunctional lateral line on expression of the social hormone *parathyroid hormone 2*

**DOI:** 10.1101/2022.05.06.490781

**Authors:** Alexandra Venuto, Cameron P. Smith, Marybelle Cameron-Pack, Timothy Erickson

## Abstract

Parathyroid hormone 2 (Pth2) is a vertebrate-specific neuropeptide whose thalamic expression is upregulated by social contact with conspecifics. However, social interactions fail to stimulate *pth2* expression in isolated zebrafish whose lateral line hair cells have been chemically ablated. These results suggest that modulation of *pth2* by social context is acutely dependent on mechanosensory information from the lateral line. However, it is unclear how a congenital loss of lateral line function influences the ability of zebrafish to interpret their social environment. In this study, we measure *pth2* levels in zebrafish mutants lacking hair cell function in either the lateral line only, or in both the inner ear and lateral line. Socially-raised lateral line mutants express lower levels of *pth2* relative to wild type siblings, but there is no further reduction when all sensory hair cells are nonfunctional. However, social isolation of hair cell mutants causes a further reduction in *pth2* expression, pointing to additional unidentified sensory cues that influence *pth2* production. Lastly, we report that social context modulates fluorescent transgenes driven by the *pth2* promoter. Altogether, these data suggest that lateral line mutants experience a form of isolation, even when raised in a social environment.

**SUMMARY STATEMENT:** Expression of the pro-social neuropeptide *pth2* is downregulated in larval zebrafish with a congenital loss of lateral line function. Thus, even in social environments, fish with compromised lateral lines may experience a form of isolation.

## INTRODUCTION

The complex relationship between the brain and social behavior has been studied across animal species for decades (Brothers, 1990; Chen and Hong, 2018; Kravitz and Huber, 2003; Robinson et al., 2008). Different social contexts can evoke a wide range of emotions, from joy and positivity (Burgdorf and Panksepp, 2006; Panksepp and Burgdorf, 2003) to fear and aggression (Amaral, 2002; Neumann et al., 2010) with a high level of plasticity (Chen and Hong, 2018; Kolb and Whishaw, 2003). In situations of social isolation, many animals exhibit anxious behaviors in addition to neurochemical changes in the brain (Amaral, 2002; Mumtaz et al., 2018; Shams et al., 2017; Weiss et al., 2004).

Social behavior is modulated by a suite of neuropeptides (Adkins-Regan, 2009; O’Connell and Hofmann, 2011; Oldfield and Hofmann, 2011; Panksepp et al., 2007; Schoofs et al., 2017). Parathyroid hormone 2 (Pth2, formerly known as tuberoinfundibular peptide of 39 residues or TIP39) is a neuropeptide expressed in the vertebrate thalamus, which is a sensory integration and relay center (Anneser et al., 2020; Bhattacharya et al., 2011; Dobolyi et al., 2012; Mueller, 2012). In rodents, the PTH2 receptor is expressed in several nuclei throughout the central nervous system (Dobolyi et al., 2012). Zebrafish have two *pth2r* paralogs, *pth2ra* and *pth2rb*, each with distinct expression patterns in the fish brain (Seitz, 2019). Activation of PTH2R increases intracellular cAMP levels and can modulate neuroendocrine activity in the hypothalamic– pituitary–adrenal (HPA) axis (Dimitrov and Usdin, 2010). PTH2-PTH2R signaling influences several behaviors, including maternal care (Cservenák et al., 2013), pain sensitivity (Dimitrov et al., 2010; Dimitrov et al., 2013; LaBuda and Usdin, 2004), and anxiety-like responses (Anneser et al., 2022; Coutellier and Usdin, 2011; LaBuda et al., 2004).

Recent work suggests that Pth2 signaling promotes social interactions between conspecifics. In zebrafish, *pth2* mutants exhibit decreased preference for conspecifics (Anneser et al., 2022), while chemogenetic activation of the Pth2-positive region of the rodent brain promotes social contact (Keller et al., 2022). Consistent with this, expression of *pth2* itself positively scales with the quantity of social interactions with conspecifics (Anneser et al., 2020; Anneser et al., 2022; Keller et al., 2022). In rodents, physical touch from conspecifics activates *Pth2* expression (Cservenák et al., 2013; Keller et al., 2022). In addition to tactile touch sensation, fish also have another form of “distant touch” through the mechanosensory lateral line (Dijkgraaf, 1963). The mechanosensory lateral line is a hair cell-based system in aquatic vertebrates that senses hydrodynamic information, such as that created by conspecifics (Butler and Maruska, 2015; Butler and Maruska, 2016a; Butler and Maruska, 2016b; Groneberg et al., 2020; Montgomery et al., 2013). In zebrafish, chemical ablation of the mechanosensory lateral line prevents the rescue of *pth2* expression by a social environment following isolation (Anneser et al., 2020). Therefore, social “touch” (direct contact in rodents and hydrodynamic communication in zebrafish) seems to be a primary sensory activator of *pth2* expression.

Despite this knowledge, there remain several unanswered questions regarding the social regulation of *pth2* expression. First, how do zebrafish interpret social context when they are born without lateral line function? Secondly, do mechanosensory hair cells in the inner ear contribute to the ability of zebrafish larvae to respond to social context? Thirdly, is *pth2* expression regulated solely via mechanosensation, or can fish use additional sensory inputs to sense the presence of conspecifics? Here, we report that the congenital loss of mechanosensory hair cell function leads to chronically depressed levels of *pth2* expression in socially-housed larval zebrafish. Interestingly, social isolation of hair cell mutants further decreases *pth2* levels, suggesting that larval fish use additional sensory information to sense conspecifics and modulate *pth2* expression. Lastly, we utilize fluorescent reporters driven by the *pth2* regulatory region to evaluate the morphology of *pth2*-expressing cells and visualize *pth2* dynamics *in vivo*. Overall, this study offers additional insight into the sensory basis for social interactions in zebrafish and introduces novel tools to track *pth2* expression in living fish.

## RESULTS

### Decreased *pth2* expression in lateral line-deficient larval zebrafish

*pth2* expression in the thalamus decreases when fish are raised in isolation but increases following a brief period of social interaction with conspecifics. However, when the mechanosensory lateral line is ablated in isolated fish prior to social exposure, *pth2* expression levels remain low (Anneser et al., 2020). This implicates the lateral line sensory system in recognizing conspecifics to stimulate *pth2* expression. To further evaluate the role of the lateral line in social recognition, we raised lateral line-deficient mutants (*lhfpl5b*^*-/-*^) and their siblings in separate social groups and detected *pth2* expression by mRNA *in situ* hybridization (ISH) at 4 days post-fertilization (dpf). Socially-raised lateral line mutants exhibit a decrease in *pth2* expression (M = 9.7, SD = 4.3) compared to wild type (WT) siblings (M = 29.9, SD = 1.7) (Fig. 1B, C), in terms of a qualitative decrease in the intensity of staining as well as a significant decrease in the number of detectable *pth2*-positive cells (Fig. 1D; One-tailed Welch’s t-test, *t* = 18.25, *P* = 1.04×10^−15^, Cohen *d* = 6.18). We also ablated lateral line hair cells in socially-raised WT larvae using repeated neomycin treatments between 3 – 4 dpf (Venuto and Erickson, 2021) (Fig. 1E, Fig. S1). Even though these larvae had social experience prior to losing lateral line input, neomycin-treated larvae exhibit a significant decrease in the number of *pth2*-positive cells compared to untreated controls (Fig. 1F; One-tailed Welch’s t-test, *t* = 6.44, *P* = 1.21×10^−7^, Cohen *d* = 3.22). These results confirm that a loss of lateral line function, whether genetically or chemically-induced, impairs the ability of *pth2* cells to respond to social context.

**Fig. 1:**
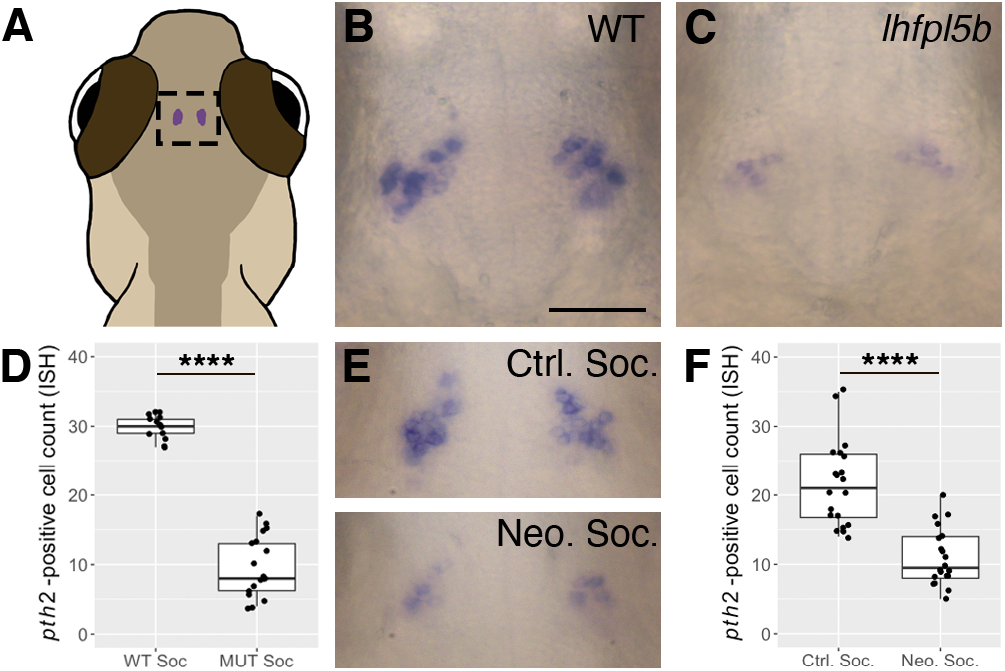
Decreased *pth2* expression in lateral line-deficient zebrafish larvae. (A) Location of *pth2*-expressing cells in the thalamic region of a larval zebrafish. (B-C) Representative images of mRNA *in situ* hybridizations (ISH) showing *pth2*-positive cells in socially-reared wild type and lateral line mutant (*lhfpl5b*^*vo35*^) larvae at 4 dpf. (D) Boxplots of *pth2*-positive cell counts from ISH images of *lhfpl5b* larvae at 4 dpf (WT = 14, MUT = 18 larvae from two replicates). (E) Representative mRNA ISH images from socially-reared control (Neo. ctrl) and neomycin-treated (Neo. trt.) larvae at 4 dpf showing reduced *pth2* expression following repeated chemical ablation of the lateral line hair cells. (F) Boxplots of *pth2*-positive cell counts from control and neomycin-treated larvae at 4 dpf (Control = 20, Treated = 20 larvae pooled from two replicates). Scale bar in B is 50 μm and applies to all images. A one-tailed Welch’s t-test was used to assess significance, **** = p < 0.0001.

### Lack of evidence for modulation of *pth2* expression by mechanosensory inputs from inner ear hair cells

Low levels of *pth2* expression remain in socially-raised lateral line mutants (Fig. 1B, C). To determine if mechanosensory input from the inner ear was responsible for this remaining *pth2* expression, we used three different types of mutations that disable hair cells of both the lateral line and inner ear to compare to the lateral line mutant (*lhfpl5b*^*-/-*^) and wild type zebrafish. Two of these mutations (*pcdh15a*^*-/-*^ and *tomt*^*-/-*^) render the mechano-electrical transduction channel nonfunctional in all hair cells, similar to how the *lhfpl5b* mutation specifically disables lateral line hair cells (Erickson et al., 2017; Erickson et al., 2020; Maeda et al., 2017). The third mutant disables the voltage-gated calcium channel in all hair cells (*cacna1da*^*-/-*^), thereby blocking stimulus-dependent glutamate release (Sheets et al., 2012; Sidi et al., 2004). Again, we measured *pth2* expression via mRNA ISH (Fig. 2B-F) as well as quantitative PCR (qPCR) at 4 dpf (Fig. 2G). Consistent with our ISH results from Fig. 1, each class of hair cell mutant exhibits a significant decrease in *pth2* expression compared to their wild type siblings when raised socially (Fig. S2; One-tailed Welch’s t-test; *lhfpl5b*: *t* = 5.929, *P* = 0.011, Cohen *d* = 4.84; *tomt*: *t* = 3.857, *P* = 0.0305, Cohen *d* = 3.15; *cacna1da*: *t* = 5.154, *P* = 0.0044, Cohen *d* = 4.21). However, the pan-hair cell mutants (*tomt, cacna1da*) did not differ in *pth2* expression levels compared to the lateral line-only mutant (*lhfpl5b*) (Fig. 2G; One-way ANOVA; Mutant: *F* = 2.425, *P* = 0.169; Wild Type: *F* = 0.791, *P* = 0.496). Because pan-hair cell mutants and lateral line-only mutants are indistinguishable in terms of *pth2* expression levels, we conclude that inner ear mechanosensory hair cells do not significantly contribute to the regulation of *pth2* expression in this context.

**Fig. 2:**
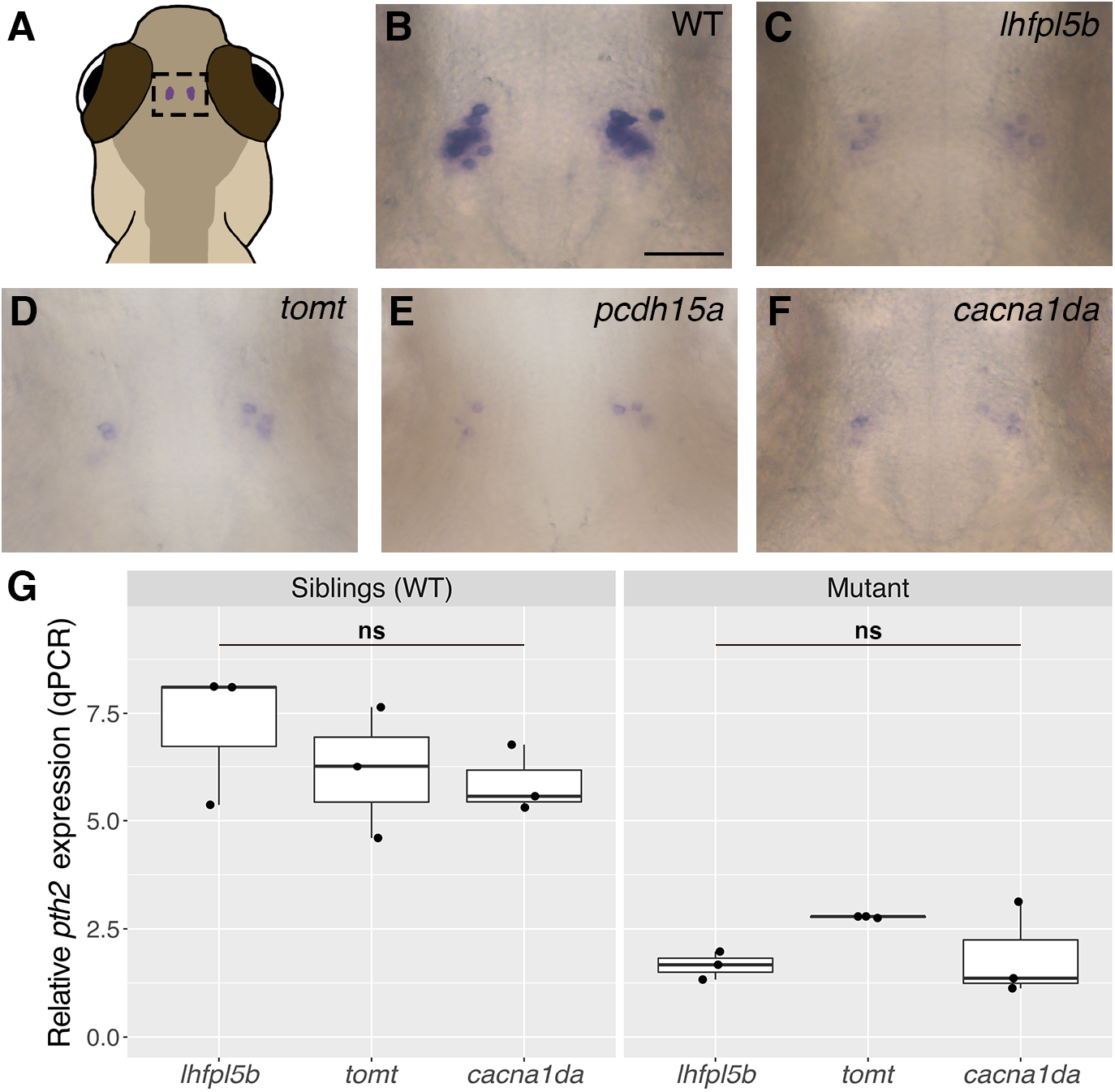
Lack of evidence for modulation of *pth2* expression by mechanosensory inputs from inner ear hair cells. (A) Location of *pth2*-expressing cells in the thalamic region of a larval zebrafish. (B-F) Representative images of mRNA *in situ* hybridizations (ISH) showing *pth2*-positive cells in socially-reared B) wild type (*n* = 14), C) *lhfpl5b*^*vo35*^ (*n* = 18), D) *tomt*^*tk256c*^(*n* = 13), E) *pcdh15a*^*th263b*^ (*n* = 13), and F) *cacna1da*^*tc323d*^ (*n* = 18) larvae at 4 dpf. Scale bar = 50 μm, applies to all images. G: Boxplots of qPCR results showing *pth2* expression in socially-reared mutants and siblings at 4 dpf. Ten larval heads were pooled per condition to make one biological replicate, and three biological replicates were used to generate the data. Two technical replicates of the qPCR experiment were conducted per biological replicate and averaged. One-way ANOVA to assess significance, ns = not significant.

### Social isolation further reduces *pth2* expression in sensory hair cell mutants

Chemosensory and visual perception of conspecifics does not significantly rescue *pth2* expression in larval zebrafish raised in isolation (Anneser et al., 2020). However, since disabling both the lateral line and the inner ear does not completely eliminate *pth2* expression, other non-hair cell-based sensory systems may contribute to the modulation of *pth2* by social context. To test this, we compared *pth2* expression in social and isolated *lhfpl5b* and *tomt* mutants by both mRNA ISH (Fig. 3A-C) and qPCR at 4 dpf (Fig. 3D). If *pth2* responds to conspecific signals solely via the lateral line, we expect to see equivalently low expression in mutants raised in social and isolated environments. However, we find this not to be the case. Social environment has a significant effect on *pth2* expression, as isolated *lhfpl5b* and *tomt* mutants express less *pth2* than social mutant siblings (Two-way ANOVA; *lhfpl5b: F* = 16.42, *P* = 0.007; *tomt*: *F* = 65.53, *P* = 0.00019). Genotype does not have a significant effect on *pth2* levels when larvae are raised in isolation, as isolated mutants and isolated wild types are statistically indistinguishable (Two-way ANOVA; *lhfpl5b: F* = 2.65, *P* = 0.154; *tomt*: *F* = 0.873, *P* = 0.386). Because social isolation further reduces *pth2* expression in lateral line mutants, we conclude that *pth2* likely responds to multisensory input during social experience.

**Fig. 3:**
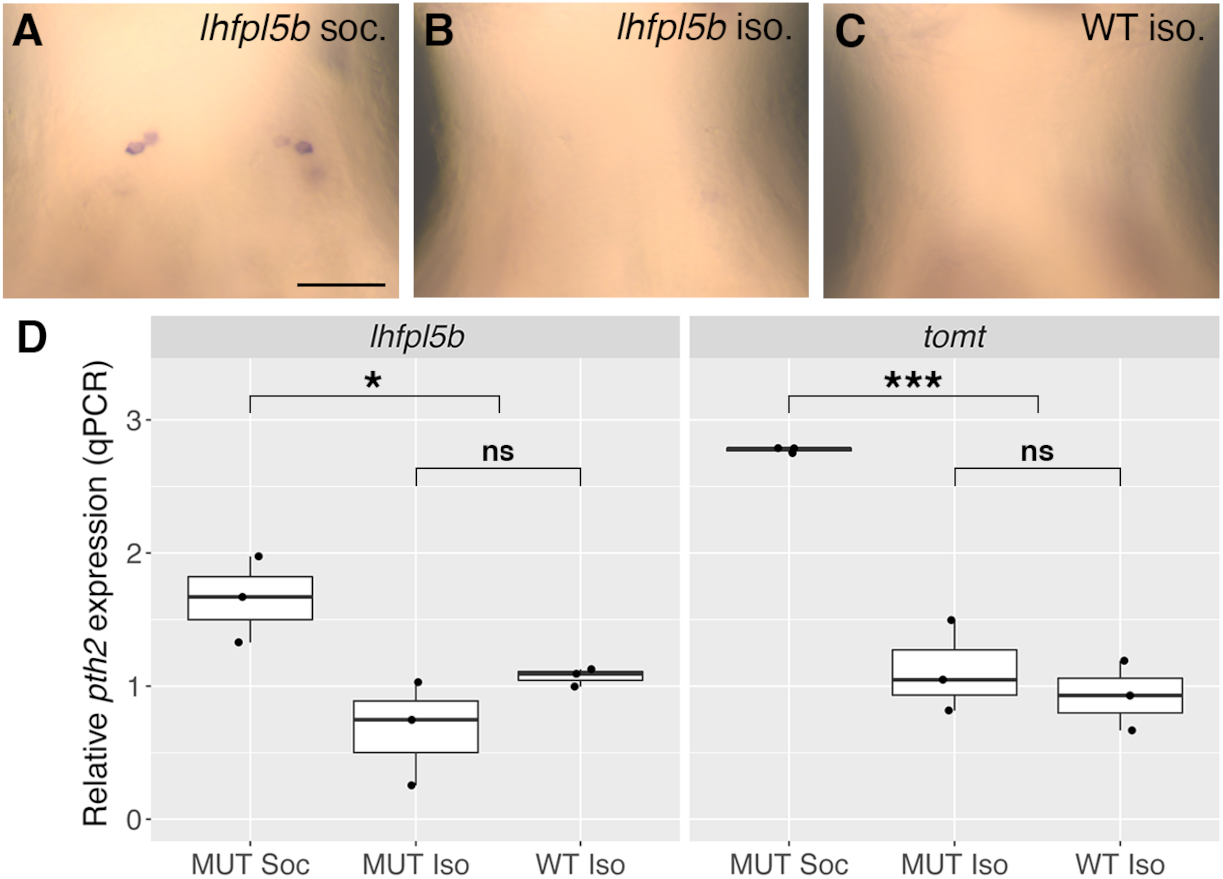
Social isolation further reduces *pth2* expression in sensory hair cell mutants. (A-C) mRNA *in situ* hybridization showing representative examples of *pth2*-positive cells in 4 dpf A) socially-reared *lhfpl5b*^*vo35*^ mutants (*n* = 48), B) isolated wild type larvae (*n* = 50), and C) isolated *lhfpl5b*^*vo35*^ mutants (*n* = 40). Scale bar = 50 μm, applies to all images. D: Boxplots of qPCR results comparing *pth2* expression in socially-reared *lhfpl5b*^*vo35*^ and *tomt*^*tk256c*^ mutants relative to isolated mutant and wild type siblings at 4 dpf. Ten larval heads were pooled per condition to make one biological replicate, and three biological replicates were used to generate the data. Two technical replicates of the qPCR experiment were conducted per biological replicate and averaged. Two-way ANOVA to assess significance, *** = p < 0.001, * = p < 0.05, ns = not significant.

### Morphology of *pth2*-expressing cells using fluorescent reporters

Fluorescent reporters provide information about the morphology and function of cells, along with the ability to quantify gene expression in living organisms (Choe et al., 2021; Soboleski et al., 2005). Therefore, we created stable lines of transgenic zebrafish expressing GFP or RFP under control of the presumptive *pth2* promoter (Fig. 4 and Movie S1). Imaging of socially-reared, transgenic larvae reveals the two bilateral cell clusters with complex projection patterns (Fig. 4A, B). The anterior ipsilateral projections partially converge into a dense neuropil in the forebrain, while other fibers project ventrally and posteriorly toward the midbrain. We also find ventral projections that cross the midline between the two cell clusters. The projection patterns of the neurites and number of cells are similar for both the GFP and RFP transgenes (Fig. S3; GFP: M = 38.5, SD = 2.8; RFP: M = 37.1, SD = 2.6; Two-tailed Welch’s t-test; *t* = 1.422, *P* = 0.166, Cohen *d* = 0.519). Combining counts from both clusters, there is an overall average of 40.3 cells per individual (SD = 2.27). The cells within each cluster exhibit a range in fluorescent intensity, and over half of the cells fall within the two lowest intensity bins (Fig 4C, D; M = 27.9, SD = 3.42).

**Fig. 4:**
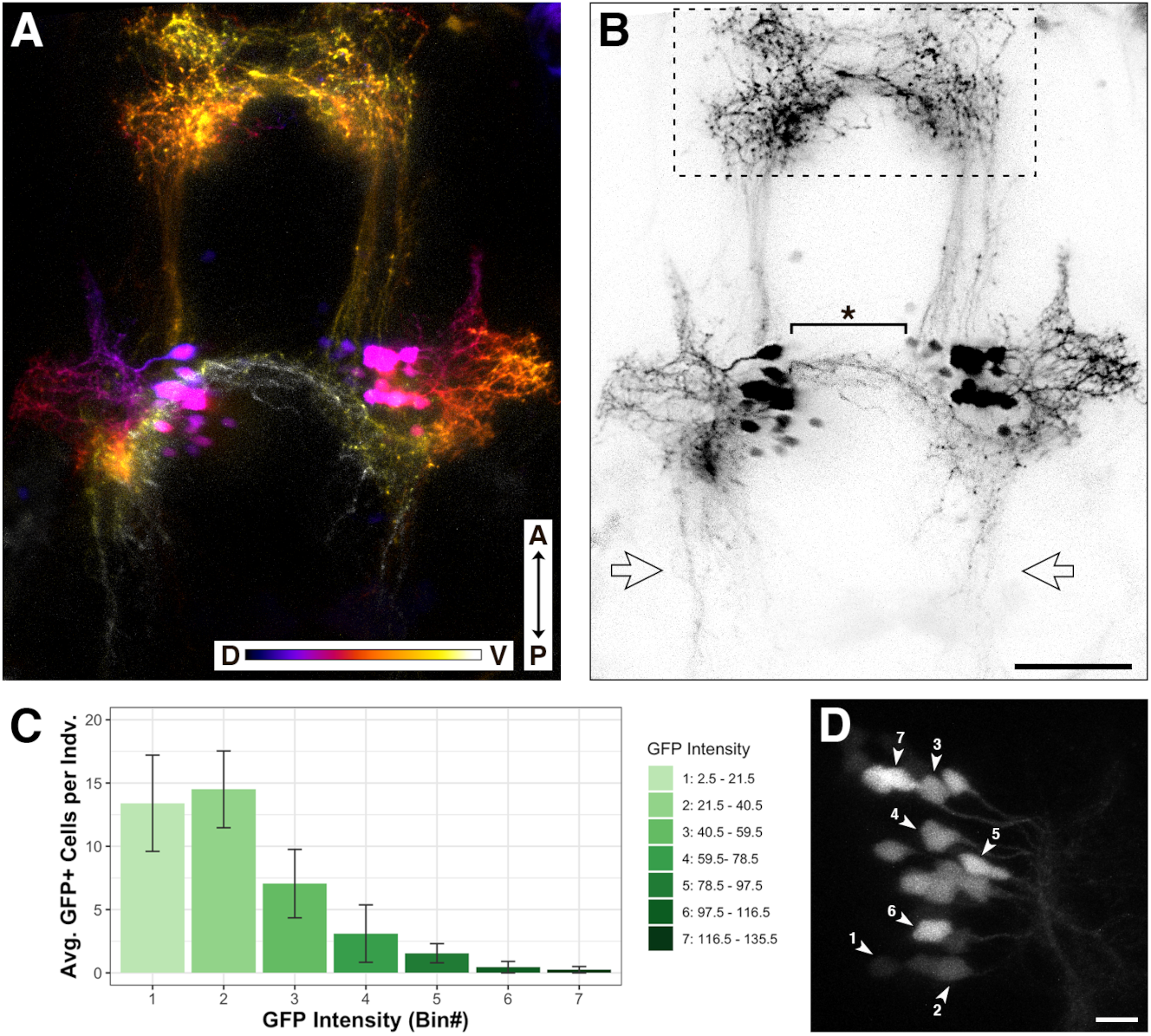
Morphology of *pth2*-expressing cells using fluorescent reporters. A: Depth-encoded representation of the *Tg(pth2:TagRFP)unb3* transgenic line at 5 dpf. The anterior-posterior (A-P) axis and dorsal-ventral (D-V) depth color scale are indicated at the bottom. Image depth is 152 μm. B: Black and white version of the image in A. From the top: the dashed box indicates the anterior neuropil, the starred bracket highlights the ventral neurites that cross the midline, and the bilateral open arrows point to posterior neurites. Scale bar in B is 50 μm, applies to both images. See Movie S1 for a 3D rendering of the depth-encoded *pth2* transgene. C: Bar graph showing binned fluorescent intensity of GFP-positive cells (n = 20 larvae from 2 different clutches at 4 dpf). D: Zoomed-in image of one cell cluster exhibiting the full range of cell fluorescent intensity, labelled according to bins outlined in the associated bar graph. Image depth is 32 μm. Scale bar is 10 μm.

### The *pth2* transgenes respond to social context and lateral line input

Next, we tested if the *pth2* transgenes respond to social context and lateral line input like endogenous *pth2* (Fig. 5). Consistent with our mRNA ISH and qPCR results, isolated *Tg(pth2:TagRFP)unb3* larvae exhibit a significant decrease in the average number of TagRFP-positive, *pth2* cells (false-colored green) (M=28.3, SD = 2.8) relative to socially-reared siblings (M = 37.3, SD = 2.6) at 4 dpf (Fig. 5A-C: One-tailed Welch’s t-test; *t* = 8.949, *P* = 5.42×10^−10^, Cohen *d* = 3.27). Similarly, social *Tg(pth2:EGFP)unb2*; *lhfpl5b*^*vo35/vo35*^ larvae have fewer fluorescent cells on average (M = 34.2, SD = 2.8) than their social, WT siblings (M = 43.2, SD = 3.9) (Fig. 5D-F; One-tailed Welch’s t-test; *t* = 6.421, *P* = 1.55×10^−6^, Cohen *d* = 2.62). To gauge whether isolation specifically diminishes expression from the *pth2*-driven transgenes, we compared the effect of social isolation on transgene expression in *Tg(pth2:TagRFP)unb3; Tg(slc6a3:EGFP)ot80* double transgenic larvae. The *slc6a3:EGFP* transgene expresses GFP under control of the dopamine transporter (DAT) promoter in several discrete regions of the zebrafish brain (Xi et al., 2011), including a population of dopaminergic cells in the pretectum lying just dorsal to the *pth2*-expressing cells. If social context specifically modulates expression from the *pth2:TagRFP* transgene, we expect that isolation will have no effect on the number of neighboring dopaminergic cells. Indeed, the number of GFP-positive, dopaminergic cells (false-colored magenta) in the pretectum do not change in response to social context (Fig. 5A-C; WT Soc: M = 44.0, SD = 4.3; WT Iso: M = 45.1, SD = 4.7; One-tailed Welch’s t-test, *t* = -0.654, *P* = 0.741, Cohen *d* = 0.239). These experiments demonstrate that the *pth2* transgenes specifically respond to both social context and lateral line input. However, we note that the effects of isolation and lateral line-deficiency on the *pth2* transgenes are not as pronounced as their effects on endogenous *pth2* mRNA expression (Figs. 1 and 2). To test whether this discrepancy is due to the high stability of green and red fluorescent proteins (Sacchetti et al., 2001; Verkhusha et al., 2003), we compared the number of GFP-positive cells at 4 dpf when we begin isolation at 1 dpf vs. 2 dpf (Fig. S4). We find that increasing the duration of isolation does not significantly change the number of GFP-positive cells (1 dpf Iso: 29.8, SD = 2.6; 2 dpf Iso: M = 29.3, SD = 3.4; Two-tailed Welch’s t-test; *t* = -0.371, *P* = 0.716, Cohen *d* = 0.166). This indicates that the discrepancy between transgenic and endogenous *pth2* expression is not due to insufficient isolation time.

**Fig. 5:**
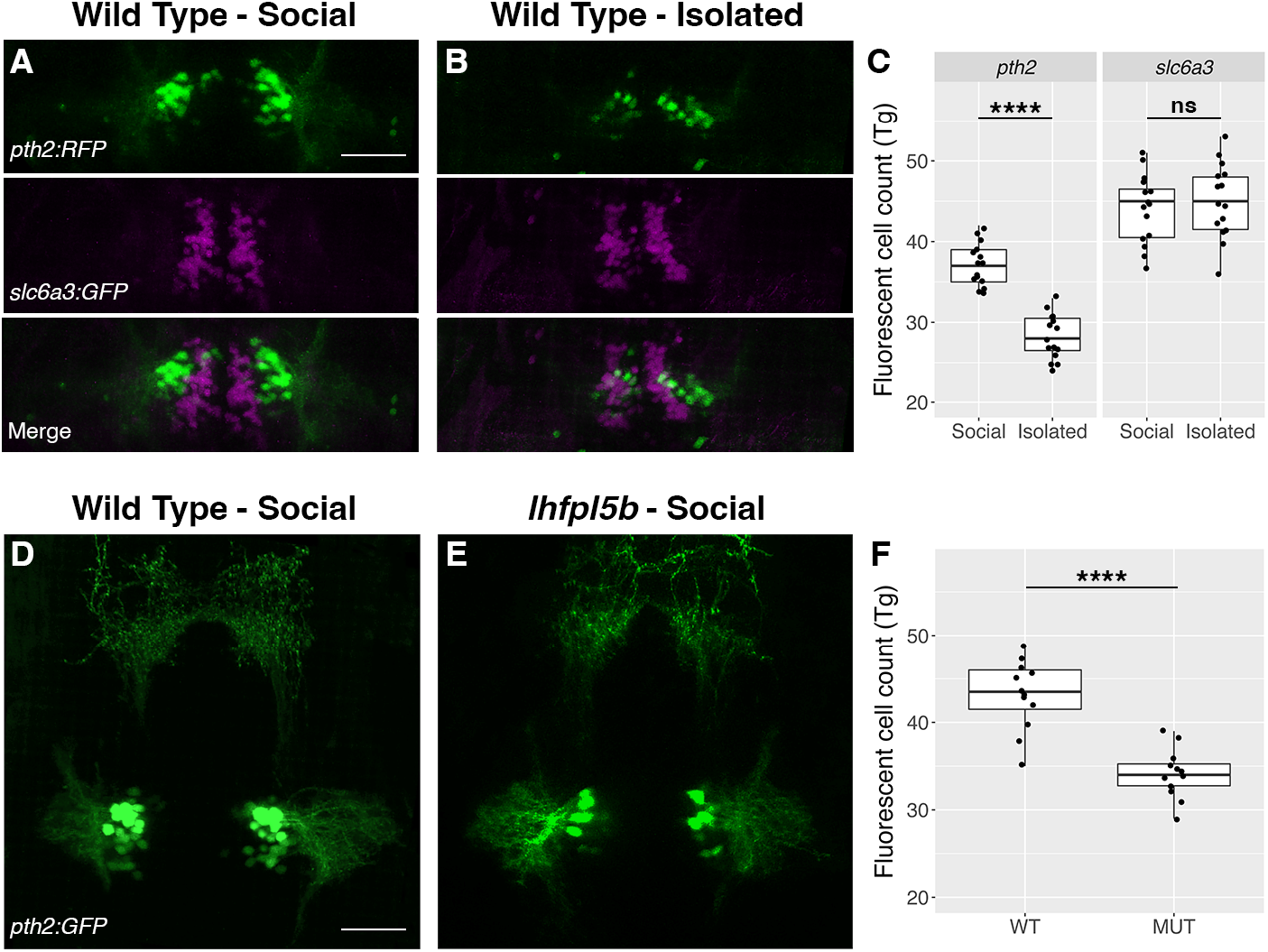
The *pth2* transgenes respond to social context and lateral line input. (A-B) Representative images of *Tg*(*pth2:TagRFP)unb3* (false-colored green) and *Tg(slc6a3:EGFP)ot80* (false-colored magenta) transgenes in socially-reared and isolated larvae at 4 dpf. Merges of the two channels show the close proximity of the *pth2* and *slc6a3-*positive cell clusters. (C) Boxplots of *pth2* and *slc6a3* cell counts (*n* = 15 per condition). (D-E) Representative images of *Tg*(*pth2:EGFP)unb2* transgene expression in socially-reared wild type and lateral line mutant (*lhfpl5b*^*vo35*^) larvae at 4 dpf. (F) Boxplots of GFP-positive cell counts in wild type (WT) and *lhfpl5b* mutant (MUT) siblings (*n* = 12 per condition). Scale bar is 50 μm. For both experiments, a one-tailed Welch’s t-test was used to assess significance, **** = p < 0.0001, ns = not significant.

## DISCUSSION

Previous work revealed that larval zebrafish use lateral line information to sense the presence of conspecifics, leading to upregulation of the social hormone Pth2. Here, we show that *pth2* levels are chronically depressed in socially-raised larvae born without lateral line function. However, social isolation further reduces *pth2* levels in lateral line mutants, suggesting that other socially-derived sensory information can modulate *pth2* expression.

### The mechanosensory lateral line mediates *pth2* expression

Anneser, et al. (2020) implicated hydrodynamic stimuli from conspecifics as a key social cue that upregulates production of the neuropeptide Pth2. Isolated larvae with chemically ablated lateral lines fail to upregulate *pth2* when returned to a social environment. In the present study, we use mutant zebrafish with a congenital loss of lateral line function to confirm that hydrodynamic information modulates *pth2* expression. Furthermore, we find that repeated chemical ablation of the lateral line also impairs the ability of social larvae to express *pth2*, even when exposed to conspecifics prior to loss of lateral line function. Our approach complements that of Anneser et al. by providing genetic evidence that *pth2* is modulated by lateral line sensory information (Fig. 1). We conclude that, in terms of Pth2 production, zebrafish born without a lateral line experience a form of isolation even when raised in a social context.

### Auditory and vestibular stimuli do not modulate *pth2* expression in larval zebrafish

Inner ear hair cells are functionally and structurally similar to lateral line hair cells, and fish use auditory and vestibular sensory information in a variety of behaviors (Bagnall and Schoppik, 2018; Kohashi et al., 2012; Nicolson et al., 1998; Privat et al., 2019). Given that some *pth2* expression remains in lateral line mutants, we examined whether sensory information from the inner ear contributes to social modulation of *pth2*. Using zebrafish with mutations that disable both inner ear and the lateral line hair cells, we find that there is no additional loss of *pth2* expression compared to the lateral line-only mutant (Fig. 2). In addition, we asked if different types of mutations to hair cells impact *pth2* expression. Three of the mutant lines used in this study, *lhfpl5b, tomt* and *pcdh15a*, render the mechanotransduction (MET) channel non-functional (Erickson et al., 2017; Erickson et al., 2020; Maeda et al., 2017), and one mutant line, *cacna1da / cav1.3a*, affects the voltage-gated calcium channel (Sheets et al., 2012; Sidi et al., 2004). All mutant alleles are recognized as nulls since mutant hair cells show little to no residual mechanosensory activity, as demonstrated by the lack of evoked spikes in the posterior lateral line ganglion, the absence of microphonic potentials and mechanotransduction currents, or the absence of MET channel-dependent FM dye labeling of hair cells (Nicolson et al., 1998; Erickson et al., 2017; Erickson et al., 2020; Sidi et al., 2004). While the MET channel and calcium channel are both necessary components for proper hair cell signal transduction, there are key differences between the types of mutations. For mutants affecting the MET channel, potassium and calcium ions are unable to enter the cell in response to stimuli, but the potential for spontaneous calcium influx and neurotransmitter release remains (Trapani and Nicolson, 2011). Whereas for the calcium channel mutation, ions enter through the MET channel but there is no evoked or spontaneous neurotransmitter release. Regardless, all mutations that disrupt hair cell function result in the same decrease of *pth2* expression in socially raised fish compared to their wild type siblings (Fig. 2). Therefore, we conclude that regulation of *pth2* by social context is primarily via the mechanosensory lateral line in larval zebrafish and that auditory and vestibular information likely does not contribute.

### Additional socially-derived sensory cues modulate *pth2* expression

Anneser et al. (2020) demonstrated that neither chemical nor visual cues from conspecifics significantly restored *pth2* expression in isolated larvae. In our study, we found that isolation further reduced *pth2* expression in lateral line mutants to the same level as isolated wild types (Fig. 3). This indicates that additional sensory modalities contribute to the regulation of *pth2* expression. Recent studies have identified thalamic neurons in the zebrafish that respond to biological motion and shown that these neurons are molecularly defined by the partially overlapping expression of *pth2* and *cortistatin* (*cort* / *sst7*) (Kappel et al., 2022; Sherman et al., 2021). It will be informative to test whether photo- or chemosensory systems can modulate *pth2* expression under sensitized circumstances in which hydrosensory information is compromised. Additionally, it is possible that tactile stimuli are contributing to *pth2* regulation in zebrafish, consistent with the effects of social touch in rodents. Future work is necessary to test this theory.

### Visualizing *pth2* expression *in vivo*

We used *pth2* promoter-driven fluorescent transgenes to visualize *pth2*-expressing cells and their projections. Consistent with the Anneser et al. (2020) whole-mount immunostains, we see Pth2-positive cells located in bilateral clusters in the thalamus that feature lateral neurites, anterior ipsilateral projections terminating as a loosely organized neuropil in the telencephalon, as well as posterior ipsilateral projections. The transgene reveals additional fibers that project ventrally from the cell bodies and cross the midline, potentially contacting the contralateral *pth2*-expressing cluster (Fig. 4 and Movie S1). Investigating the morphology of these cells will help us understand the cellular targets of Pth2 signaling and how activity of the two clusters is coordinated. We note an excess of putative *pth2*-expressing cells in the transgenics compared to our ISH experiments and the immunostain counts of Anneser et al. (2020). This discrepancy could be due, in part, to the increased sensitivity of using confocal microscopy to detect *pth2*-expressing cells in a live transgenic animal as opposed to imaging fixed tissue stained for mRNA or protein. Consistent with this idea, we find that the majority of GFP-positive cells in socially-reared *Tg(pth2:EGFP)* larvae exhibit low fluorescence intensity (Fig. 4C, D). If these dim transgenic cells represent neurons that express *pth2* at a level below the detection limit of mRNA ISH or immunostain, this may account for the excess cell counts in our transgenics.

We also tested the *pth2:EGFP* and *pth2:TagRFP* transgenic lines as a proxy for *pth2* expression *in vivo* and found that both lateral line mutants and isolated wild type larvae exhibit fewer fluorescent cells than socially raised siblings (Fig. 5). In contrast, the number of dopaminergic cells located dorsally to the *pth2* clusters exhibit no change to social context. Dopamine has been implicated in social interactions (Haehnel-Taguchi et al., 2018; Saif et al., 2013; Shams et al., 2018), so we used the nearby clusters of GFP-labeled dopaminergic cells in the *slc6a3:EGFP* transgenic line to test the specificity of the response of the *pth2* transgene to social context. Our results indicate that the social modulation of *pth2* is specific and distinct from the dopaminergic system in this brain region of larval zebrafish.

While our results with the *pth2* transgenes agree with our *in situ* hybridization and qPCR results (Figs. 1-3), we note that the effects of isolation and lateral line deficiency on transgene expression are not as prominent as when assaying for endogenous *pth2* mRNA and protein. As mentioned above, the enhanced detectability of genetically-encoded reporters in live larvae may account for some of the residual fluorescence that remains following social isolation of transgenics. It could also be that the post-transcriptional and / or post-translational stability of the fluorescent transgenes differ from endogenous *pth2* mRNA and protein stability. We tested whether the long half-life of GFP accounts for the persistence of GFP-positive cells in isolated *Tg(pth2:EGFP)* larvae but found that extending the duration of isolation from 48 to 72 hours did not significantly affect the number of fluorescent cells (Fig. S4). Lastly, it is possible that additional regulatory elements in the *pth2* promoter or 3’ UTR, along with the use of a destabilized fluorescent protein, are required to mimic the endogenous regulation of *pth2*. Future work will strive to distinguish between these possibilities and to optimize the transgene for more accurate reporting of *pth2* dynamics in vivo.

Overall, our results confirm that *pth2* expression is modulated by the mechanosensory lateral line (Fig. 1). We also show that the lateral line is the only hair-cell based sensory organ regulating *pth2* expression (Fig. 2). However, due to expression levels being slightly higher in social lateral line mutants compared to the same mutants or wild type larvae raised in isolation, we conclude that there must be unidentified sensory system(s) that contribute to the modulation of *pth2* (Fig. 3). Finally, we created transgenic lines to visualize the morphology of *pth2*+ cells *in vivo* (Fig. 4), and report that the transgenes respond to social context (Fig. 5). These transgenic lines can be used in future studies to elucidate the role of *pth2* in zebrafish. Taken together, this work suggests that zebrafish without a functioning lateral line experience a form of isolation, even in a social context.

## MATERIALS AND METHODS

### Ethics statement

Animal research complied with guidelines stipulated by the Institutional Animal Care and Use Committees at East Carolina University (Greenville, NC, United States) and at University of New Brunswick (Fredericton, New Brunswick, Canada). Zebrafish (*Danio rerio*) were maintained and bred using standard procedures (Westerfield, 2000). All experiments used larvae at 1–5 days post-fertilization (dpf), which are of indeterminate sex at this stage.

### Mutant and transgenic fish lines

The following previously described zebrafish mutant and transgenic lines were used in this study: *lhfpl5b*^*vo35*^, *pcdh15a*^*th263b*^, *tomt*^*tk256c*^, *cacna1da*^*tc323d*^, and *Tg(slc6a3:EGFP)ot80* (Erickson et al., 2017; Erickson et al., 2020; Maeda et al., 2017; Sidi et al., 2004; Xi et al., 2011). To make the *Tg(pth2:EGFP)unb2* and *Tg*(*pth2:TagRFP)unb3* transgenic lines, 2067 base pairs immediately upstream of the *pth2* coding region (including the 5’ UTR and first intron) were PCR amplified with primers containing attB4-B1 recombination sites (*pth2*-specific sequence underlined: Fwd: GGGGACAACTTTGTATAGAAAAGTTGGTAAAGGACACTCTTTGAGATCCT, Rvs: GGGGACTGCTTTTTTGTACAAACTTGCCTGCAGATGAATAAGTTGAATAATACA. The PCR product was cloned into the Gateway pDONR-P4-P1R entry vector by a standard BP clonase reaction and a Multisite LR reaction was performed to create *pth2:EGFP-pA, cryaa:EGFP* and *pth2:tagRFP-pA, cryaa:mCherry* plasmids. pDestTol2pACryGFP and pDESTtol2pACrymCherry plasmids were gifts from Joachim Berger & Peter Currie (Addgene plasmids # 64022, 64023). The expression constructs were co-injected with *tol2* transposase mRNA into 1-cell zebrafish embryos (Kwan et al., 2007). F0 founders were outcrossed to produce stable F1 transgenics. At a minimum, progeny from F2 transgenics were used in this study.

### Mutant and social isolation experiments for *in situ* hybridization, qPCR, and transgenic lines

Zebrafish larvae were sorted for *lhfpl5b*^*vo35*^, *pcdh15a*^*th263b*^, *tomt*^*tk256c*^ homozygotes at 2 dpf with FM1-43 dye labelling for social isolation experiments. *cacna1da*^*tc323d*^ homozygotes were identified behaviorally by the “circler” phenotype at 4 dpf (Nicolson, et al., 1998), as they cannot be identified by differential FM1-43 dye labeling and were not used in isolation experiments. *Tg*(*pth2:EGFP)* and *Tg*(*pth2:TagRFP)* transgenics were identified by their GFP or mCherry lens markers, respectively. *Tg(slc6a3:EGFP)* transgenics were identified by GFP expression in the brain. Identified larvae were divided into social groups of thirty, while another thirty larvae were placed in isolation, except for *cacna1da*^*tc323d*^ larvae which were only raised socially. Both social and isolated larvae were housed in 6-well tissue culture plates containing 5 ml of E3 embryo media. For the social condition, all 30 larvae were placed in one well with 5 ml of embryo media. For the isolated condition, one larva was placed per well with 5 ml of embryo media. Strips of paper were placed between each well to prevent visual access between wells. Larvae intended for mRNA *in situ* hybridization (ISH) were grown in E3 with 200 μM N-Phenylthiourea (PTU) (Alfa Aesar) to prevent pigment formation. Each group was incubated at 28.5ºC with a 14L:10D photoperiod from 2 to 4 dpf. At 4 dpf, larvae were either fixed for *in situ* hybridization, processed for qPCR, or imaged to visualize the transgene.

### Repeated neomycin treatments

The repeated neomycin treatments were performed essentially as previously described (Venuto & Erickson, 2021). At 24 hpf, WT embryos were divided into groups of 30 and grown continuously in the dark at 28.5ºC in media supplemented with 200 μM PTU to prevent pigment formation. Larvae were manually dechorionated at 2 dpf. Starting at 9:00 AM, 3 dpf larvae were exposed to either 0 or 50 μM neomycin sulfate hydrate (Alfa Aesar) for 30 minutes at 28.5ºC every 12 hours for a total of three treatments. At noon on 4 dpf, treated and untreated groups of larvae were anesthetized with 0.016% MS-222 and fixed in 4% PFA for mRNA ISH. The experiments were repeated three times, each with two technical replicates for the control and treatment groups. Three hours after treatment 1, ablation of lateral line hair cells was assessed by incubating 3 dpf larvae in 130 μM DASPEI (2-[4-(dimethylamino)styryl]-1-ethylpyridinium iodide) for 10 minutes. Following three washes with E3, larvae were anesthetized with 0.016% MS-222, positioned laterally on a depression slide in 1.2% low melting point agarose, and imaged with a Basler Ace 5.1 MP color camera on a Zeiss SV-11 microscope using an X-Cite 120Q illuminator and wideband GFP filter.

### mRNA in situ hybridization

The *pth2* in situ probe sequence was amplified using the Invitrogen SuperScript IV One-Step RT-PCR kit (ThermoFisher) using the following forward and reverse primers: F: CATTGCATGGACGATTTACG; R: TGCCATGTCATTCAAAATCC. The RT-PCR product was purified by gel extraction and propagated in the pCR4-TOPO vector. The pCR4-*pth2* plasmid was linearized with NotI and the antisense DIG-labeled ISH probe was synthesized using the T3 RNA polymerase promoter. Probe synthesis and in situ hybridization were performed essentially as described (Erickson et al., 2010). Larvae were imaged at 20X magnification using either an Olympus BH-2 microscope outfitted with a Jenoptik Gryphax Arktur camera or a Nikon Eclipse 50i microscope with a Basler Ace 5.1 MP color camera. ISH cell counts were conducted in a non-blinded manner by manually focusing through a mounted larvae prior to capturing a static image.

### RNA isolation and real time quantitative PCR (qPCR)

For RNA isolation, ten larval heads were used per biological replicate. Samples were collected on ice, placed in 1.7 mL RNase-free microcentrifuge tubes containing 200 μl RNAzol (Molecular Research Center, Inc., OH. Catalog: RN 190) and homogenized immediately. This was repeated for a total of three biological replicates. Total RNA was extracted from homogenized solutions according to the manufacturer’s protocol. For each sample, cDNAs was synthesized using 1 μg total RNA and a high-capacity cDNA Reverse Transcription kit (Thermo Fisher Scientific, Waltham, Massachusetts, USA, Catalog# 4368814) following the manufacturer’s instructions.

The levels of *pth2* transcripts were determined by quantitative real-time PCR (qPCR) using SYBR green dye (Invitrogen) and a CFX Connect real-time thermal cycler (Bio-Rad Laboratories, Hercules, California, USA). The qPCR reaction was conducted with initial denaturation at 95 °C for 2 min, followed by 30 cycles of 30 s denaturation at 95 °C, 30 s annealing at 57 °C, and 30 s extension at 72 °C using the specific primers (Forward: CCACGCAACACACAGTCAAG, Reverse: GCAAGTTACTTTGCAGAGGTC) and GoTaq G2 DNA polymerase (Promega, Madison, Wisconsin, USA). Each PCR mixture (15 μl) consisted of 7.795 μl DNase free water, 3 μl 5X GoTaq buffer, 1.5 μl 25 mM MgCl2, 0.3 μl 10 mM dNTP mix, 0.15 ml 10μM forward or reverse primer, 2 μl cDNA, 0.03 μl 100X SYBR green dye (final concentration 0.2X), and 0.075 μl Taq. Two technical replicates per biological replicate were averaged together for analysis. Absolute values (copies/μg total RNA) were determined using Ct values of samples and a standard curve generated from serial known concentrations of plasmid containing the target region of *pth2*. Relative expression was found by calculating the fold change compared to the mean value of the isolated, wild type condition. The efficiency of the PCR and authentic PCR products was further confirmed by analyses of melting curve and gel electrophoresis.

### Transgenic imaging and quantification

Live zebrafish larvae were anesthetized with 0.016% MS-222 in E3 and mounted dorsally on a depression slide in 1.2% low melting point agarose in E3. EGFP was imaged using a Zeiss LSM800 laser-scanning confocal with a 40x water objective and Zeiss Zen software. To quantify results, cell counts were performed using Fiji/ImageJ image processing software (Schindelin et al., 2012). Cells were counted in a non-blinded manner by manually going through the z-stacks. The cell count process was repeated twice at a minimum to ensure accuracy. Images shown are maximum intensity projections of z-stacks. For the fluorescent intensity experiment, the intensity of each cell was collected from a sum slices z-projection in Fiji/ImageJ and the integrated density values were used for analysis. Depth encoded 2D and 3D projections were created in Fiji/ImageJ using the *Z-stack Depth Color Code 0.0.2* plugin. Figures were assembled in Adobe Photoshop and equally adjusted for brightness and contrast among corresponding treatment groups.

### Statistical tests

All graphs and statistical tests were done using R (R Core Team, 2021). For *in* situ hybridization cell counts, a one-tailed Welch’s t-test was used (two biological replicates, *n* = 7-9 larvae per condition per replicate). For the qPCR experiment comparing the various hair cell mutants raised socially (Fig. 2), a one-way ANOVA was conducted (three biological replicates, *n* = 10 per condition per replicate). For the same experiment comparing wild type versus mutant larvae raised socially, a one-tailed Welch’s t-test was used to compare within mutant lines. For the qPCR experiment comparing genotypes and social status (Fig. 3), a two-way ANOVA was conducted (three biological replicates, *n* = 10 per condition per replicate). For the transgenic cell counts, a one-tailed Welch’s t-test was used (three biological replicates, *n* = 5 per condition per replicate), except for the 1 dpf versus 2 dpf and GFP versus RFP cell counts where we used two-tailed Welch’s t-test to test significance. A p-value of less than 0.05 was considered significant. The more stringent one-tailed t-tests were used due to only having an expectation for a decrease in *pth2* expression following isolation or loss of lateral line sensory information. Sample sizes were determined by the number of fish used in the experiment and no samples were excluded from the analysis. Pre-determined criteria to exclude samples included if the fish was not touch-responsive or displayed obvious developmental abnormalities.

## ACKNOWLEDGMENTS

The authors would like to thank Dr. Yong Zhu (ECU) for reagents and assistance with qPCR experiments, Dr. Les Cwynar (UNB) for use of his microscope, and Dr. Bryan Crawford (UNB) for reagents and helpful comments on the manuscript. We also thank members of the Erickson and Issa labs for their assistance with zebrafish husbandry.

## DATA AVAILABILITY

The datasets generated for this study are available on request to the corresponding author.

## SUPPLEMENTAL FIGURES AND LEGENDS

**Fig. S1:**
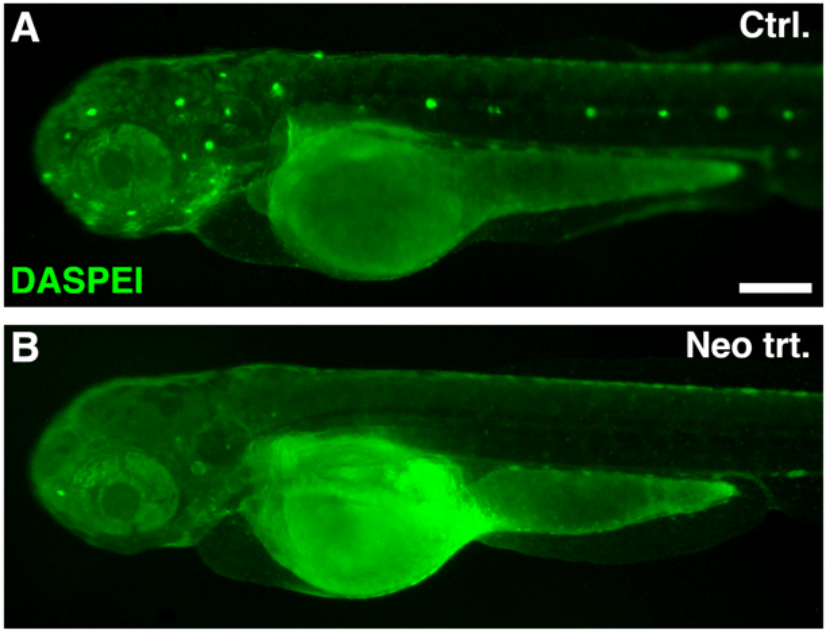
Vital dye staining of lateral line neuromasts in control and neomycin-treated larvae. Representative 3 dpf larvae from control (A) and neomycin-treated (B) groups stained with DASPEI three hours following a 30-minute treatment with either 0 μM or 50 μM neomycin. Note the near absence of punctate neuromast staining in B, indicating successful ablation of the lateral line hair cells. Scale bar = 200 μm.

**Fig. S2:**
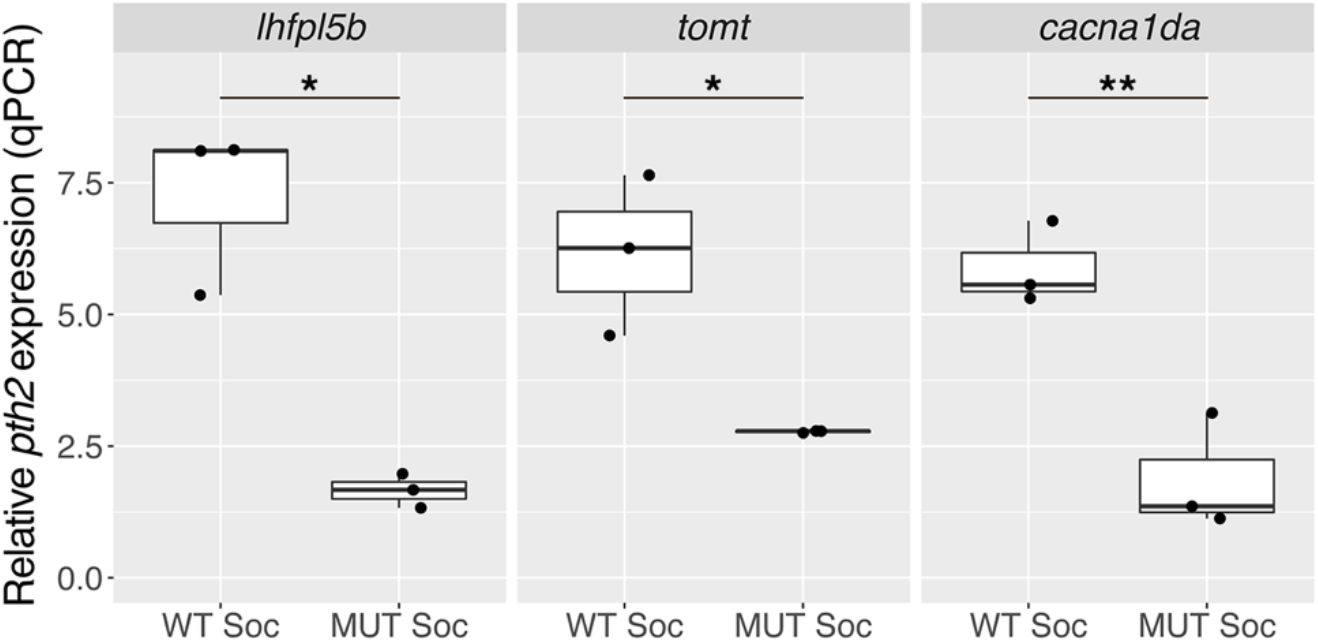
Boxplots of qPCR results showing *pth2* expression in socially-raised hair cell mutants and wild type siblings at 4 dpf. Ten larval heads were pooled per condition to make one biological replicate, and three biological replicates were used to generate the data. Two technical replicates of the qPCR experiment were conducted per biological replicate and averaged. A one-tailed Welch’s t-test was used to assess significance, * = p < 0.05, ** = p < 0.01.

**Fig. S3:**
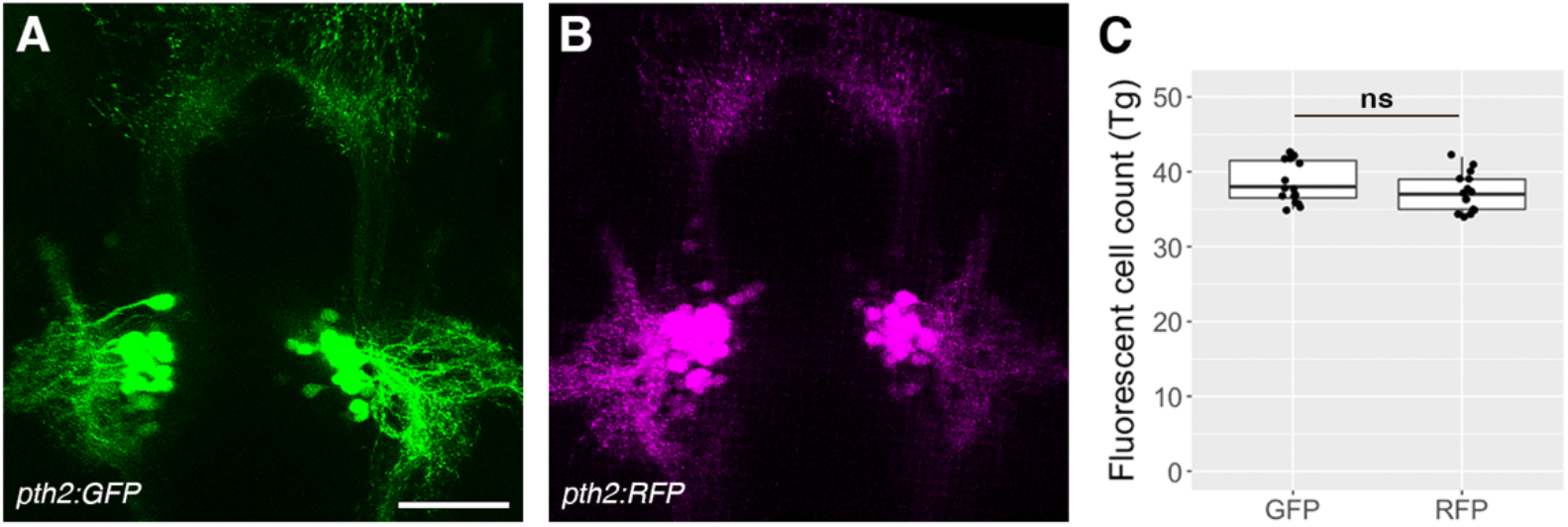
*Tg*(*pth2:EGFP)unb2* and *Tg*(*pth2:TagRFP)unb3* transgenes exhibit similar morphology and number of cells. (A-B) Representative images of socially-reared 4 dpf larva possessing either the A) *Tg(pth2:EGFP)unb2* transgene (green) or B) *Tg(pth2:TagRFP)unb3* transgene (magenta). (C) Boxplot of GFP and RFP cell counts (*n* = 15 larvae per condition). Scale bar is 50 μm. A two-tailed Welch’s t-test was used to assess significance, ns = not significant.

**Fig. S4:**
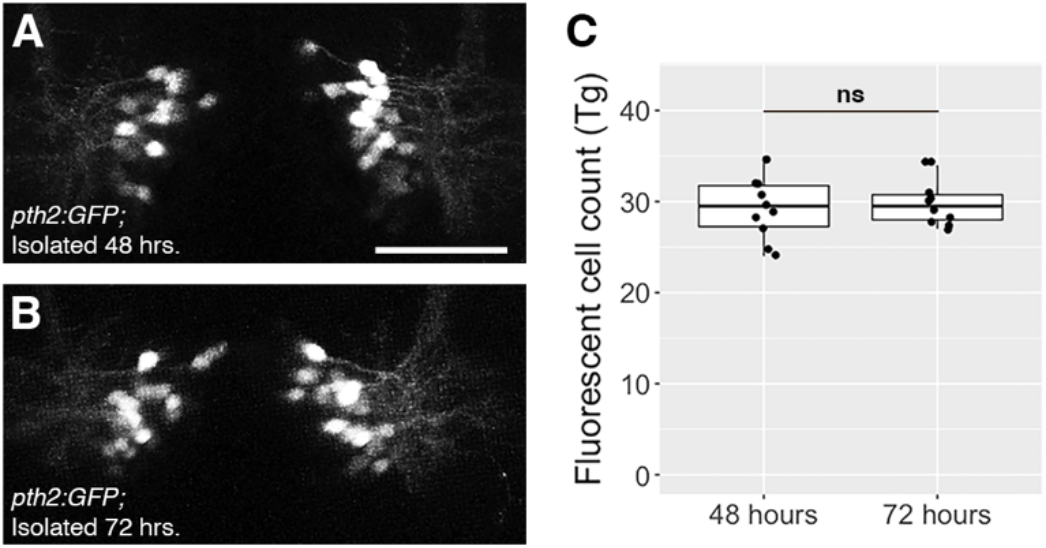
Increasing the isolation period from 48 to 72 hours does not affect the number of GFP-positive cells in *pth2:GFP* transgenics. (A-B) Representative images of 4 dpf *Tg*(*pth2:EGFP)unb2* larva raised in isolation for A) 48 hours or B) 72 hours. (C) Boxplots of GFP-positive cell counts for *Tg(pth2:EGFP)* larvae raised in isolation for 48 or 72 hours (*n* = 10 per condition). Scale bar is 50 μm. A two-tailed Welch’s t-test was used to assess significance, ns = not significant.

**Movie S1:** Related to Fig. 4A - Movie showing a 3D representation of cell morphology and neurite projections in a *Tg(pth2:TagRFP)unb3* zebrafish at 5 dpf. Projection depth is 152 μm.

